# FETCH: A platform for high-throughput quantification of gap junction hemichannel docking

**DOI:** 10.1101/2021.06.07.447352

**Authors:** Elizabeth Ransey, Kirill Chesnov, Nenad Bursac, Kafui Dzirasa

**Author notes:** **Correspondence should be sent to:** Kafui Dzirasa, M.D. Ph.D., Dept. of Psychiatry and Behavioral Sciences, Duke University Medical Center, 421 Bryan Research Building, Box 3209, Durham, NC 27710, USA, Twitter: @KafuiDzirasa.

## Abstract

Gap junctions are membrane spanning channels that connect the cytoplasm of apposed cells, allowing for the passage of small molecules and ions. They are formed by the connexin (Cx) family of proteins which assemble into hexameric hemichannels on each cell and dock to create gap junctional channels between two cells. Despite importance of various Cx isoforms in human physiology and disease, available tools for screening and discriminating their interactions such as hemichannel compatibility, docking and permeability are limited. Here, we developed FETCH (<u>f</u>low <u>e</u>nabled <u>t</u>racking of <u>c</u>onnexosomes in <u>H</u>EK cells), a method which utilizes the generation of annular gap junctions (connexosomes) as downstream indicators of hemichannel compatibility for intercellular docking. First, we show that fluorescent connexosomes create a cellular phenotype that is detectable by flow cytometry analysis. We then show that FETCH identifies homotypic and heterotypic docking of many single isoform connexin hemichannels. Finally, we demonstrate that FETCH captures the impact of disease-relevant connexin protein mutations on gap junction formation. Thus, we establish a new flow cytometry-based method that is amenable to the high-throughput classification of gap junction hemichannel docking.

## INTRODUCTION

Gap junctions (GJs) are multimeric, transmembrane channels that play crucial roles in tissue homeostasis, cellular signaling, and the propagation of electrical current by enabling direct intercellular communication between adjacent, apposed cells via ionic and small molecule exchange (reviewed in [1]). GJs are composed of connexin (Cx) proteins, which oligomerize into hexameric hemichannels [2, 3] and dock with compatible hemichannels of adjacent cells at the plasma membrane to form an intercellular pore connecting the two cells [4]. Connexins comprise a family of integral membrane proteins consisting of 21 unique isoforms in humans [5, 6], with at least one isoform expressed in virtually every tissue and major organ in the body [7, 8] resulting in distinct and overlapping isoform-specific expression patterns. Routinely, single cell-types simultaneously express two or more connexin isoforms, creating both opportunities for functional redundancy and complex, highly-regulated tissue communication (reviewed in [9]).

Connexins play significant roles in the normal physiology of the tissues in which they are expressed, thus, mutations in any of the Cx genes, can lead to pathological changes. For example, at least 100 mutations of the Connexin26 (Cx26) encoding gene (GJB2) accounts for half of all worldwide cases of congenital sensorineural deafness [10, 11]; Charcot-Marie-Tooth, a motor and sensory neurodegenerative disorder is associated with mutations of the Connexin32 (Cx32) gene (GJB1) [12] and at least 73 different mutations of the most ubiquitously expressing connexin, Connexin43 (Cx43) gene (GJA1) are associated with Oculodentodigital dysplasia (ODDD) [13], a rare, developmental disorder resulting in numerous morphological anomalies and neurological symptoms (reviewed in [14]). Additionally, aberrant expression levels or regulation of Cx43, the primary connexin expressed in the heart, are associated with cardiac arrythmias in the context of myocardial ischemia [15].

Connexin proteins share a conserved topology consisting of intracellular amino- and carboxy- termini, a cytoplasmic loop, 4 transmembrane (TM) helices, and 2 extracellular loops (ELs). Post-translational modification (e.g. phosphorylation) of the Cx C-terminus regulates transport to and from the plasma membrane [16] and the N-terminus contribute to channel gating [17]; motifs in and near the transmembrane domains affect connexin oligomerization and GJ or hemichannel conductance [18]; and the ELs primarily function to impart hemichannel docking specificity (reviewed in [19]), though EL1 can also contribute to channel properties including permeability and conductance [17–24]. Thus, the primary differentiating features of connexins are trafficking and assembly, oligomerization specificity, docking specificity and permeability or conductance.

The presence of multiple unique Cx isoforms throughout the body diversifies GJ intercellular communication. For example, many Cxs form homotypic GJs (two hemichannels of one isoform, e.g. Cx36/Cx36 GJs between two neurons [25], but some isoforms can also form heterotypic channels (two hemichannels of different isoforms, e.g. Cx46/Cx50 GJs in the lens [26] with compatibility being dictated by motifs in the ELs. Frequently, because each isoform has slightly variable permeability and conductivity, heterotypic channels exhibit a preferred ionic directional flow (rectification). Additionally, within cells in which more than one Cx is expressed, multiple isoforms may be mixed into single hemichannels, called hetero-oligomerized or heteromeric hemichannels (e.g. cardiac Cx43/Cx45 hemichannels [27], with compatibility dictated by motifs within and adjacent to TM regions [19, 28].

Another important feature of GJ intercellular communication centers on the lifetime of connexin proteins at the plasma membrane being relatively short with half-lives of ~1-5 hrs [29–32]. Given the critical utility of Cxs throughout the body, this short half-life suggests that Cxs are maintained in a constant flux of biosynthesis and degradation, which would allow a cell to rapidly up- or down- regulate GJ intercellular communication in response to physiological needs. Importantly, once hemichannels dock, they are virtually inseparable under physiological conditions [33, 34], thus, down-regulation of GJ intercellular communication is achieved by functional GJ internalization. Specifically, GJs are turned over from the plasma membrane via a unique clathrin/dynamin-dependent internalization process [35–37] that results in the internalization of portions of entire GJ plaques in the form of double-bilayer vesicular structures, termed annular gap junctions or, more recently, connexosomes [38–40]. Connexosomes contain fully-docked gap junctions that are either degraded [35] or recycled and transported back to the plasma membrane [41–43].

Despite numerous, broad implications of connexin isoforms in human physiology and disease, available tools for discerning key attributes such as hemichannel docking and permeability in vitro are limited. The most readily available methods for detecting activity often rely on dye transfer (e.g., GAP-FRAP, scrape loading and microinjection) which depend on the permeability of channels to small molecule dyes such as calcein, carboxyfluorescein, Lucifer yellow, and ethidium bromide, and use the transfer of these dyes between adjacent cells as indicators of functional gap junctions [44]. However, very few dyes have been established as gap junction permeant and such dyes are generally connexin isoform, isoform species and pH specific [45]. The highest precision and gold-standard method for evaluating connexin activity is whole-cell, dual patch clamp measurement [46, 47], which can quantitatively characterize the electrical coupling (conductance) between two cells mediated by gap junctions with high sensitivity. However, this is a very low-throughput, labor intensive and expensive technique. Thus, a higher throughput, universal method for detection/determination of connexin interactions would contribute more comprehensive understanding of connexin biology.

Here, we developed a fluorescence-based, flow cytometry method for the evaluation of gap junction formation. Our method, termed <u>f</u>low cytometry <u>e</u>nabled <u>t</u>racking of <u>c</u>onnexosomes in <u>H</u>EK cells (FETCH) is easy, accessible, fast and scalable for high-throughput analysis. Specifically, FETCH exploits the generation of fluorescent connexosomes between transfected HEK cell populations expressing distinct connexins. As connexosomes are comprised of docked hemichannels from two different cells, we determined that we could evaluate the docking capability of different connexin isoforms or mutant proteins by assessing the presence or absence of formed connexosomes, a phenotype that is discernable by flow cytometry (Figure 1).

**Figure 1.**
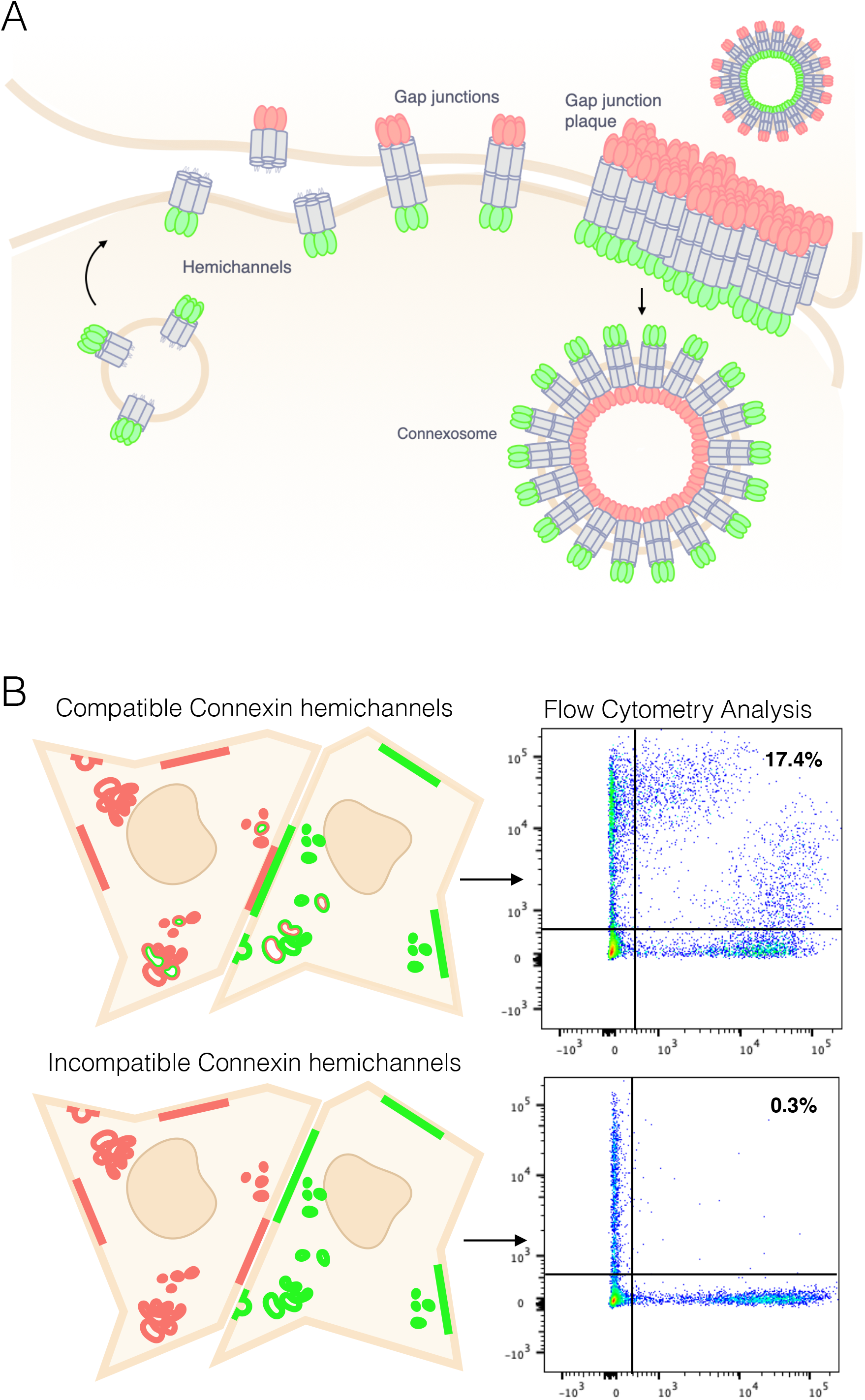
FETCH: Flow Enabled Tracking of Connexosomes in HEK cells. (A) Schematic of connexin protein life cycle. Connexin proteins assemble into hexameric hemichannels that localize to the plasma membrane of individual cells. Hemichannels diffuse laterally and dock to hemichannels on the plasma membrane of an apposed cell, forming a functional gap junction. Gap junctions concentrate at gap junction plaques which may be comprised of tens to tens of thousands of gap junctions. Cellular down-regulation of gap junction intercellular communication results in the internalization of central portions of gap junction plaques as double-bi-layer vesicular structures called connexosomes. Connexosomes are comprised of fully-docked gap junctions, thus differential fluorescent labeling of counterpart hemichannels results in dual-labeled connexosomes. (B) Schematic representing expected results of flow cytometry analysis for different combinations of connexin proteins. Paired connexin hemichannels that successfully dock are expected to elicit a strong flow cytometry phenotype representative of two-color connexosomes generation (double positive cells). Paired connexin hemichannels that fail to dock, do not generate connexosomes and, thus, do not generate a corresponding flow cytometry indicative of double positive cells.

## RESULTS

### 1. Dual fluorescently-labeled connexosomes are detectable by flow cytometry

Connexosomes are cytoplasmic, double-bilayer vesicular structures comprised of tens to thousands of fully-docked gap junction channels that have been internalized from gap junction plaques [40]. As connexosomes are the direct products of gap junction turnover, we reasoned that they could be used as indicators of gap junction docking and, by extension, as indicators of the compatibility of connexin hemichannels. Connexin43 (Cx43) is one of the best characterized connexin isoforms, and C-terminally fused fluorescent Cx43 constructs were among the first used to observe and characterize the dynamic features of connexosomes including internalization rate, vesicular sizes and cellular fates [35, 40, 48, 49]. Thus, we chose to generate two-color, fluorescent connexosomes of Cx43 and determine the feasibility of high-throughput connexosome identification in cells.

Though previous studies have evaluated endogenous connexosomes in native tissues and cells, such as membrana granulosa cells of the rabbit ovarian follicle [50–53], guinea pig epithelia [54] or the adrenocortical tumor cell line (SW-13)[55–57], or as products of exogenous construct expression in HeLa cells [35, 48, 49] or normal rat kidney (NRK) cells [40], we selected the HEK 293FT cell line (ThermoFisher) for our studies, as we observed substantially higher protein expression and larger gap junction plaque sizes as compared to HeLa and other HEK derivative cell lines (Supp. Figure 1). To generate two-color connexosomes and characterize them using microscopy, we transfected separate populations of cells with one of two fluorescently-labeled, C-terminally fused Cx43 constructs - Cx43-mEmerald or Cx43-mApple. Following an initial incubation, transfected cells were trypsinized, co-plated and incubated together for ~18 hours prior to analysis (Figure 2A). Similar to reports from other cell lines, transfection of HEK 293FTs with fluorescently-tagged Cx43 constructs resulted in the generation of connexosomes. Specifically, when we co-plated Cx43-mEmerald and Cx43-mApple expressing cells, we observed junctional plaques with dual-fluorescent co-localized invaginations, suggestive of active connexosome formation, and dual-fluorescent vesicular structures consistent with connexosomes (Figure 2B) [35, 43, 58]. In contrast, co-plated cells expressing cytoplasmic fluorescent proteins, did not generate fluorescently-labeled vesicles or plaque structures (Figure 2B).

**Figure 2.**
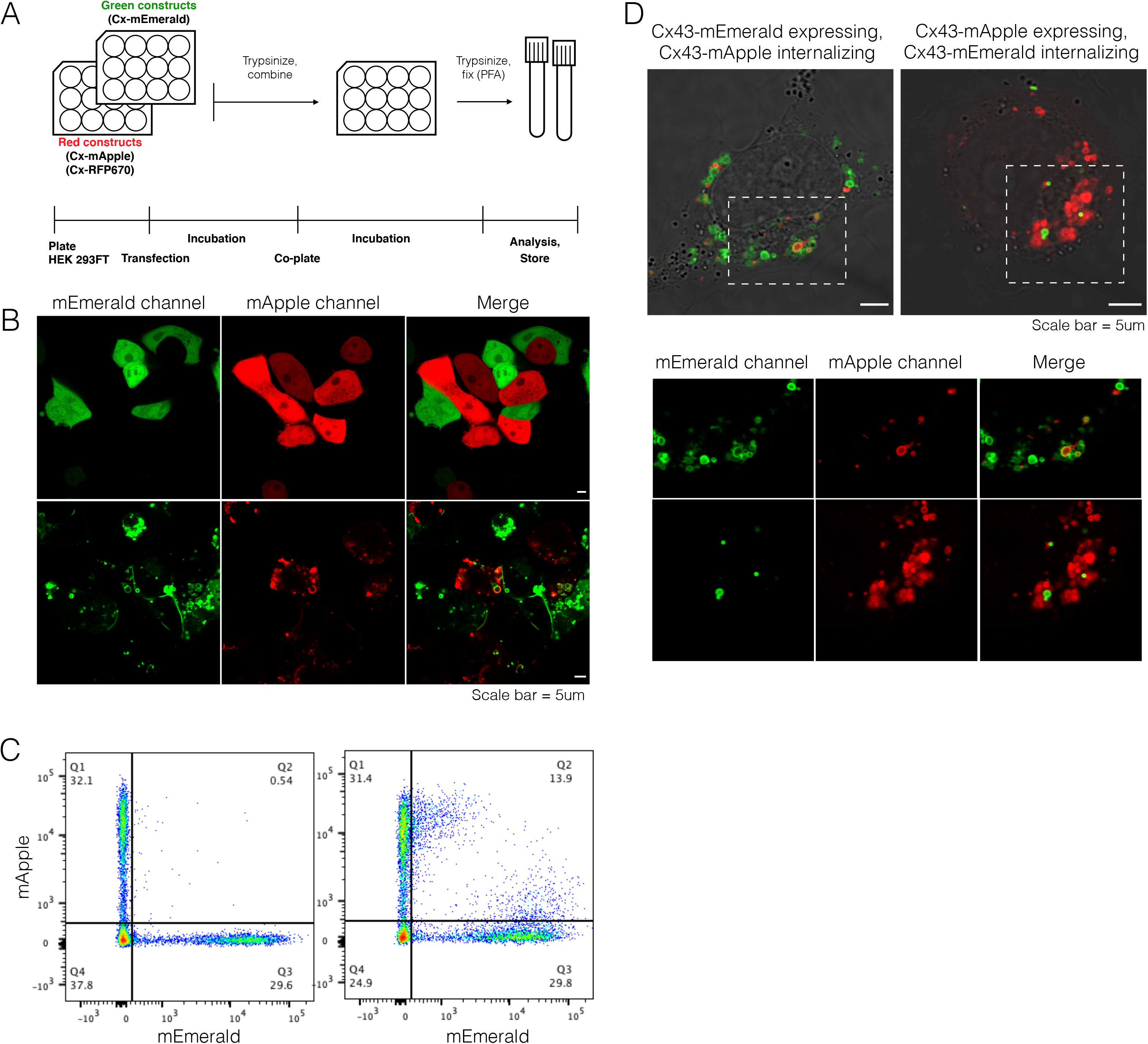
Fluorescence exchange phenotype is mediated by connexin proteins. (A) Schematic of FETCH experimental set-up involving the co-plating and incubation of individually transfected cell populations. (B) Confocal images of co-plated cytoplasmic-fluor and Cx43-fluor expressing HEK 293FT cells. (C) Flow cytometry profiles of co-plated cytoplasmic-FP and Cx43-FP cells. Fluorescence exchange can be observed as the cells that display both fluorescent proteins, despite only expressing one construct; Q2. (D) Cells sorted from either side of the double-positive quadrant of the co-plated Cx43-mEmerald/Cx43-mApple sample (i.e., higher expressing in mEmerald, higher expressing in mApple). Insets show internalized vesicular structures of double-positive sorted cells. Dual-labeled structures are consistent with connexosomes.

Flow cytometry generates multi-wavelength fluorescence profiles of individual cells within a given population, thus, we reasoned this analysis could be used to quantify fluorescence exchange between cells mediated by connexosomes in a high-throughput fashion. We termed this novel approach, flow enabled tracking of connexosomes in HEK 293FT cells, or FETCH. For FETCH analyses, we generated two-color connexosomes as described previously, but instead of imaging, we trypsinized and fixed cells in suspension, for evaluation by flow cytometry. As a control, samples of co-plated cells expressing cytoplasmic mEmerald and mApple were analyzed as well.

Upon generation of cellular fluorescence profiles by flow cytometry, we noticed a significant phenotypic difference between samples that generated connexosomes (i.e., Cx43-fluorescent protein), and those that did not (i.e., cytoplasmic fluorescent protein). For both samples, subpopulations of untransfected cells, mApple-labeled cells, and mEmerald-labeled cells were clearly identifiable (Figure 2C: Q4, Q1 and Q3, respectively). However, for the Cx43-mEmerald/Cx43-mApple sample, a significant proportion of cells exhibited both green and red fluorescence (Figure 2C, right panel, Q2). This unique double-positive or dual-fluorescent phenotype was characterized by symmetrical population shifts from the single-expressing quadrants. Specifically, cells highly expressing Cx43-mEmerald gained Cx43-mApple fluorescence and cells highly expressing Cx43-mApple gained Cx43-mEmerald fluorescence. Importantly, acquisition of the secondary fluorescent construct was commensurate with but not equal to the primary fluorescent construct. These observations are consistent with fluorescence exchange mediated by connexosomes.

To confirm the relationship between the fluorescence exchange phenotype of the Cx43-mEmerald/Cx43-mApple sample and connexosome formation, we used fluorescence-activated cell sorting (FACS) to collect cells from each subpopulation of the +/+ quadrant for microscopy analysis. Confocal imaging of the FACS sorted double-positive cells revealed dual-labeled vesicles consistent with connexosomes (Figure 2D). Moreover, the configuration of fluorescent labels of the connexosomes was such that the expressed, primary construct labels the outer surface of the connexosome and the acquired, internalized construct is on the interior. We observed this pattern for both Cx43-mEmerald and Cx43-mApple expressing cells. Though post-sorted cells showed significant amounts of vesicular structures that were single labeled (likely representing transport vesicles and internalized hemi-plaques), we did not observe any single-labeled vesicles of secondary fluorescent constructs. This supports our assertion that the dual-labeled vesicles are connexosomes and that connexosomes generate the observed fluorescence phenotype. Overall, these results demonstrate that FETCH can be used to capture the fluorescence exchange between cells, mediated by connexosomes and indicative of gap junction docking.

### 2. FETCH is broadly applicable for the evaluation of diverse Connexin protein docking interactions

The docking principles that generate annular gap junctions are common to all connexin proteins, thus, we reasoned that connexin isoforms with documented participation in homotypic gap junctions would be amenable to FETCH analysis. To standardize the quantification of fluorescence exchange we established a metric, the FETCH score, as the proportion of the +/+ quadrant cells, adjusted for total fluorescent events, minimizing the influence of transfection efficiency variability on results. We then generated known positive and negative FETCH score distributions representative of non-docking combinations for statistical comparison. To generate a positive distribution, we evaluated samples of homotypic pairings of Cx43-FPs with varied DNA transfection quantities and co-plated population densities to capture physical condition variabilities (i.e., small volume pipetting variance during transfection and co-plating). Additionally, we evaluated homotypic pairings of Connexin36(Cx36)-FPs, a neuronal connexin isoform that forms homotypic gap junctions [25, 59] (Figure 3A, Supp. Figure 2B). The negative distribution was comprised of aforementioned samples of co-plated cytoplasmic fluorescent protein (FP) expressing cells (i.e., mEmerald with mRFP670) and samples of co-plated cells expressing Cx43-FP and cytoplasmic FP (i.e., Cx43-GFP with cytoplasmic RFP). Additionally, we evaluated samples of non-docking combinations of different Cx isoforms. Specifically, co-plated cells expressing Cx43-FP with cells expressing Cx36-FP (in heterotypic combination) and cells expressing Cx23-FP (homotypically) (Figure 3A, Supp. Figure 2B). Though Cx36 and Cx43 individually form homotypic gap junctions in the mammalian nervous system, they have not been reported to form heterotypic channels and Cx23 fails to form docked gap junctions due to a missing EL disulfide bond [60].

**Figure 3.**
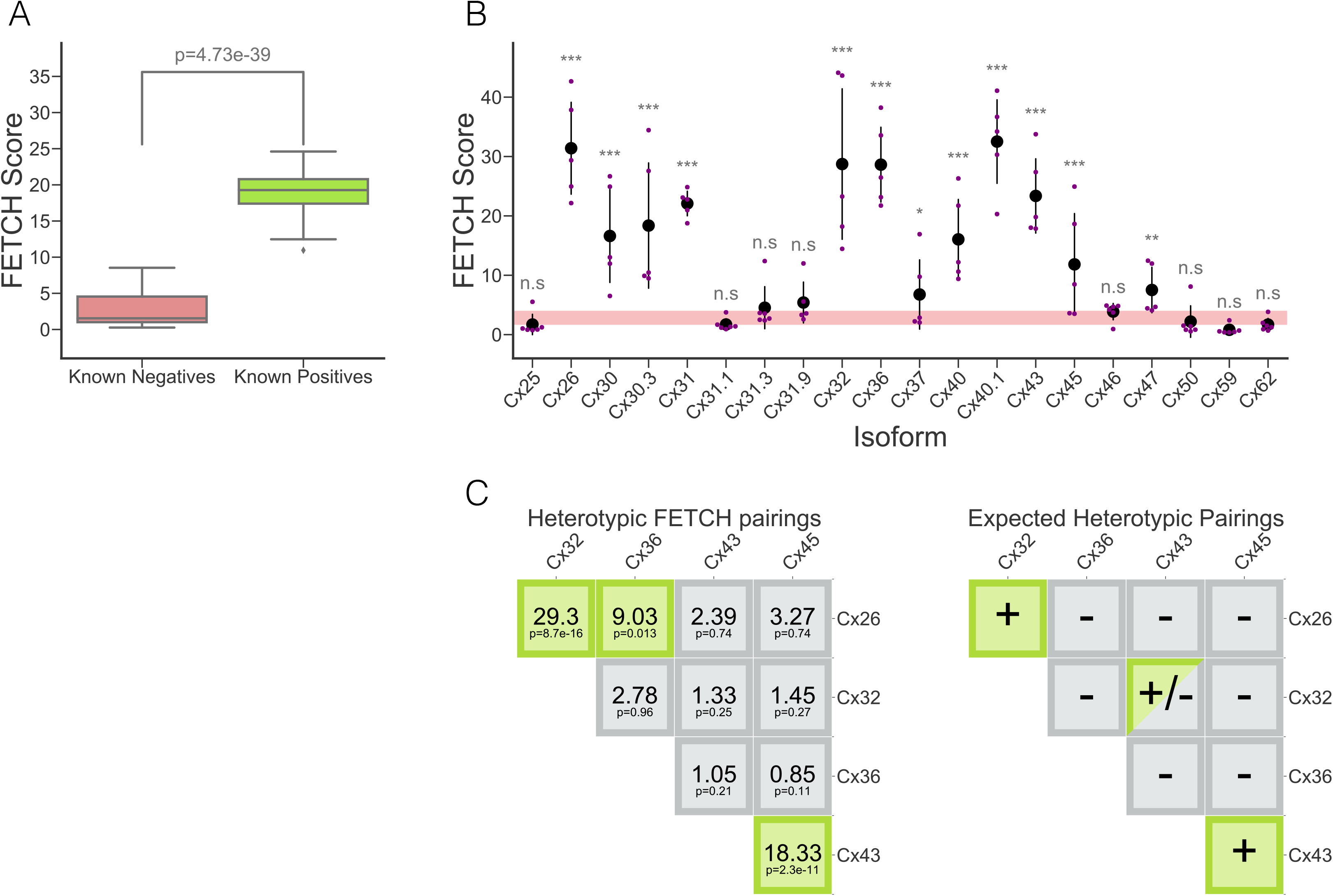
FETCH can be used to evaluate homotypic and heterotypic docking of multiple connexin isoforms. (A) FETCH score distributions of known positive and known negative connexin protein combinations. The FETCH score is defined as the proportion of Q2, double-positive cells over all fluorescent cells (Q2/Q1+Q2+Q3). (B) Homotypic FETCH scores of all human connexin isoforms. (C) Fluorescence profiles of selected isoforms (i.e., Cx26, Cx32, Cx36 and Cx45) in both homotypic FETCH and heterotypic FETCH, paired with Cx43. (D) FETCH scores of all paired combinations for Cx26, Cx32, Cx36, Cx43 and Cx45.

To evaluate the docking ability of other Cx family members using FETCH, we generated mEmerald and RFP670 C-terminal fusion constructs of the 18 remaining human connexin isoforms and completed homotypic FETCH analyses. Using an independent two-sample t-test with FDR correction, we found that FETCH scores for many Cx isoforms were significantly higher than the distribution for negative docking conditions, indicating that the majority of Cx isoforms are amenable to FETCH analysis. We observed variability in these FETCH scores, likely reflecting isoform specific characteristics (e.g., expression level, trafficking, stabilization, and turnover rate). Cx25, Cx31.1, Cx31.3, Cx31.9, Cx46, Cx50, Cx59, and Cx62 failed to generate a significant homotypic FETCH score consistent with docking (Figure 3B).

Next, to validate the specificity of FETCH in capturing compatible docking interactions, we tested select Cx isoforms for heterotypic interactions. Connexin isoforms are classified by sequence homology into 5 groups (alpha (A), beta (B), gamma (C), delta (D), epsilon (E)) [61]. Some connexin isoforms can form heterotypic (two-type) channels, frequently with other isoforms within the same homology class, and with some exceptions, isoforms in different classes are not as widely compatible for heterotypic docking (reviewed in [19, 62]). Thus, we completed heterotypic FETCH evaluation with well-characterized representatives of each major class of connexin. Specifically, in addition to Cx43, we chose Cx26, Cx32, Cx45, and Cx36 as representatives to evaluate heterotypic, combinatorial pairings. Importantly, each of these representatives exhibited significantly positive FETCH scores in homotypic evaluations.

Both Cx26 and Cx32 have previously been reported to fail to form functional heterotypic gap junctions with Cx43 in Xenopus laevis oocyte experiments [47]. On the other hand, Cx45 is frequently co-expressed in the same tissues as Cx43 and has been shown to form Cx43/Cx45 heterotypic channels [63].

When we completed heterotypic FETCH on all combinations of the subset of isoforms, we observed that most cross-class heterotypic connexin combinations yielded negative FETCH scores, reflecting docking incompatibility (Figure 3C). Using independent two-sample t-tests with FDR correction, we evaluated the FETCH scores to determine statistical significance. Consistent with expectations, FETCH analysis revealed that Cx26, Cx32, and Cx36 failed to dock with Cx43, while Cx45 showed a positive FETCH score when paired with Cx43 (n=7, z=18.33, p=2.3 × 10^−11^; Figure 3C).

In addition to the Cx43 /Cx45 pairing, we observed a positive FETCH score when pairing Cx26 with Cx32 (n=6, z=29.30, p=8.7x 10^−16^)– which belong to the same class, have homologous extracellular loop docking motifs, and have also been reported to form functional heterotypic gap junctions [23, 61, 64] (Figure 3C). Interestingly, we observed a positive FETCH score when pairing Cx26 and Cx36 (n=5, z=9.03, p=0.013), a hitherto unreported pairing that may, nonetheless, represent a newly discovered functional compatibility. These results provide strong evidence that the FETCH method captures physiological gap junction formation and may be applied to broader evaluations of other connexin isoforms in diverse and uncharacterized combinations.

### 3. FETCH captures functional docking of disease relevant connexin mutant proteins

Cx43 is the most ubiquitously expressed connexin in the body with critical, tissue-specific functions in nearly 40 tissues and organs including the heart, brain, liver and skin. Unsurprisingly, mutations in the GJA1 gene, which encodes Cx43, often result in numerous, systemic physiological perturbations. The majority of Cx43 disease-associated mutations result in the complex developmental disorder, Oculodentodigital dysplasia (ODDD) [13, 65], which is characterized by congenital craniofacial, limb and digital deformities, in addition to teeth and eye anomalies and neurological dysfunction (reviewed in [14]). To further expand upon the capabilities of the FETCH method in identifying connexin docking interactions, we selected 10 single amino acid mutations of Cx43 that are associated with the syndromic disease to evaluate for homotypic and heterotypic (against WT-Cx43) docking.

Of the 10 selected mutations associated with ODDD, 2 were localized to the intracellular N-terminus or loop (S18P and I130T), 2 were localized to transmembrane helices (A40V and V216L), 4 were localized to extracellular loop 1 (N55D, Q58H, P59A and S69Y) and 2 were localized to extracellular loop 2 (H194P and R202H) (Figure 4A). All 10 mutations are autosomal dominant and represent a range of observed channel perturbations when expressed in cultured cells and/or in mouse models. For example, the I130T and R202H mutations have been documented to form channels with reduced conduction [66, 67], whereas S18P, A40V, P59A and V216L mutations have been observed to result in a total loss of channel function (reviewed in [13]).

**Figure 4.**
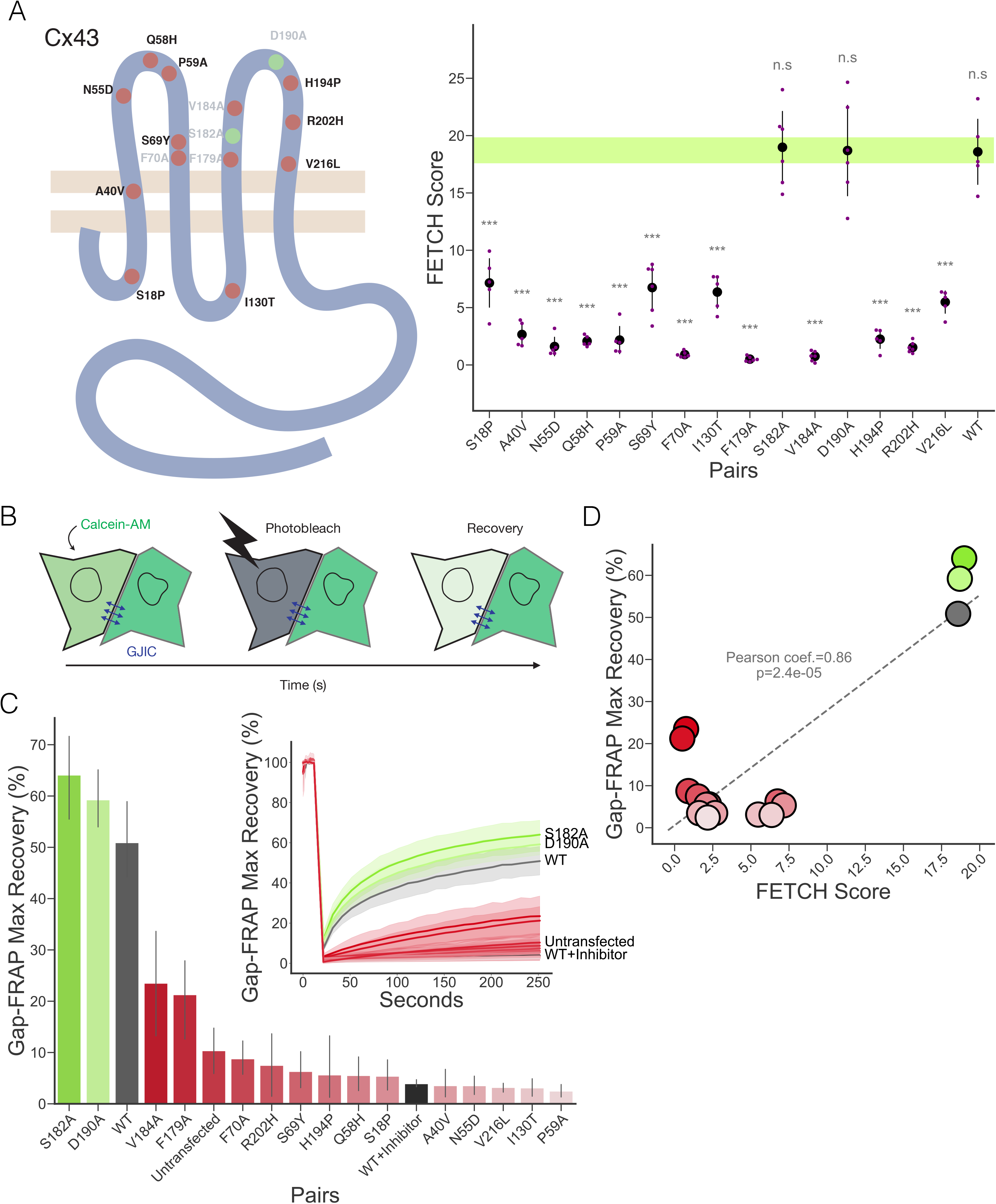
FETCH can be used to evaluate docking ability of connexin mutant proteins. (A) Schematic representation and FETCH scores of 10 Oculodentodigital dysplasia (ODDD)-associated Cx43 mutations (in black text) and 5 random alanine mutations (in gray text). Mutation colors reflect docking (green dots) and non-docking (red dots) as determined by statistical analysis of FETCH scores (B) Schematic of GAP-FRAP (fluorescence recovery after photobleaching). calcein-AM cell permeant dye is added to confluent cells transiently transfected with Cx43-WT and Cx43-mutant constructs. Upon permeating cells, calcein-AM is cleaved by amino-esterases, making it cell-impermeant; residual dye is removed via repetitive washes. Individual cells containing calcein are bleached using a 488nm laser, and the recovery of cellular fluorescence over time, mediated by gap junctions, is recorded. (C) GAP-FRAP analysis of ODDD-associated and random Cx43 mutant proteins. Bar graph depicts maximum fluorescence recovery reached at the end of the recovery time interval (4 mins). Inset shows full initial, bleach and recovery traces for each protein, in addition to corresponding 95% confidence intervals. Mutation colors are carried over from the associated FETCH analysis for each mutation as docking (green bars and traces) and non-docking (red bars and traces); Cx43-WT and Cx43-WT+Inhibitor (100uM carbenoxolone) are depicted as black.

**Figure 5.**
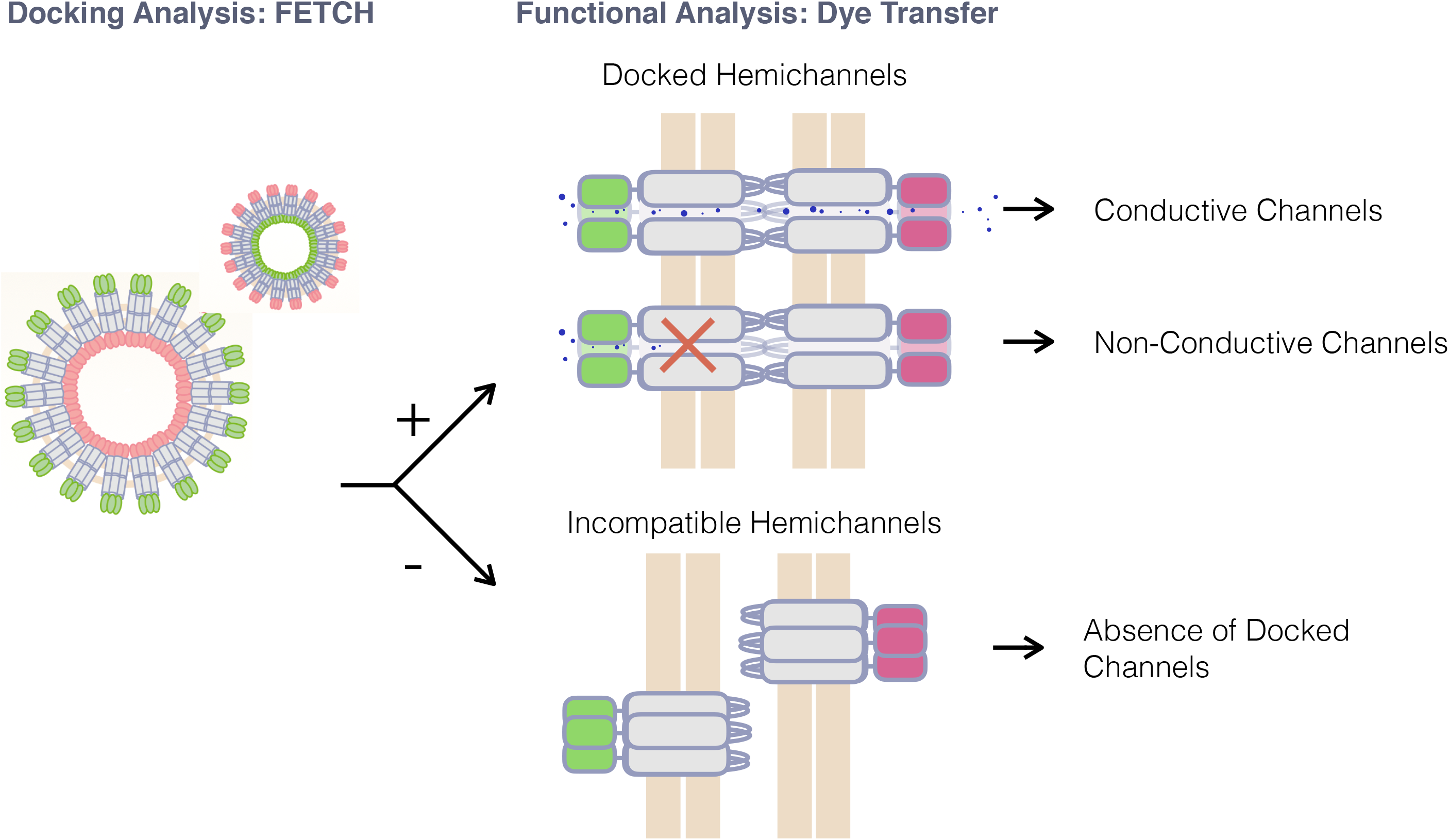
Summary of FETCH method contribution to the evaluation of connexin protein interactions. Using FETCH docking analysis coupled with a functional assay such as dye transfer can help distinguish incompatible hemichannels and non-functional, docked channels.

Since our FETCH score reflects hemichannel docking, and thus, all of the ‘upstream’ processes underlying GJ formation (i.e., Cx expression levels, trafficking, stabilization and turnover rate), we expected disease associated Cx43 mutations would exhibit diminished FETCH scores as compared to the WT-Cx43 isoform. After generating a positive control distribution for homotypic Cx43 by co-plating Cx43-FP constructs in conditions of increasing DNA transfection quantities and increasing co-plated densities (n=33), we evaluated the mutant proteins in homotypic FETCH experiments. Upon analysis, Cx43 mutant proteins demonstrated lower homotypic FETCH scores than WT-Cx43 (Figure 4A).

Since connexin hemichannel docking is a prerequisite for GJ-mediated exchange of small molecules and ions between cells, we reasoned that mutations that decrease the FETCH scores would also diminish intercellular permeability. A standard method for evaluating GJ dependent intercellular communication is fluorescence recovery after photo-bleaching (FRAP), which relies on the permeability of Cx43 gap junctions to a tracer dye (calcein-AM) as an indicator of functional channels [27, 68, 69] (Figure 4B). Thus, we evaluated mutant proteins using FRAP to determine if functional channels were being formed, despite FETCH scores suggestive of diminished docking.

Unsurprisingly, all selected Cx43 ODDD-associated mutant proteins showed significantly impaired dye transfer in FRAP experiments (Figure 4C). Thus, negative FETCH scores correlated with poor gap junction functionality as determined by FRAP. Because we were additionally interested in identifying a correlation between positive FETCH scores and FRAP-determined functionality of mutant proteins, we made random alanine scanning mutations of the Cx43 ELs at positions not explicitly characterized as disease associated. Specifically, we generated 5 constructs: F70A, F179A, S182A, V184A and D190A (Figure 4A). Upon FETCH evaluation, we found that two mutations that retained their ability to homotypically dock – S182A and D190A. Follow-up FRAP analysis demonstrated that random Cx43 mutants with positive homotypic FETCH scores were able to mediate dye transfer in FRAP assays, comparable to Cx43-WT (Figure 4C). Finally, we show that FETCH scores correlate with FRAP-determined functionality (Figure 4D). Combined, these results show that FETCH can be used to detect dysfunctional GJ intercellular communication and can characterize connexin mutant proteins for gap junction docking competency – a potential component to functional perturbation in pathological phenotypes.

## DISCUSSION

Here we describe FETCH, a novel method that employs flow cytometry to evaluate a natural product of gap junction turnover (connexosomes) as indicators of gap junction formation, a prerequisite for GJ communication. While other methods primarily quantify GJ conductive function in individual cell pairs or individual cells within a cell network, FETCH captures hemichannel docking across a population of cells. By specifically evaluating hemichannel docking compatibility, FETCH can differentiate between two potential sources of GJ non-functionality: channels that are innately non-conductive and channels that fail to form, by precisely identifying the latter. This distinction is critical for the dissection of complex mechanisms underlying both normal connexin physiology and connexin-associated pathologies. To date, most studies have evaluated Cx43 (and other Cx isoform) mutations at the level of alterations in protein expression level, localization and conduction between cells. However, sufficient expression and plasma membrane localization do not guarantee docking – a requirement for intercellular communication. Thus, discerning mutations that exclusively affect functional conduction from mutants that fail to form docked gap junctions has been a challenge. Though our small selection here did not identify Cx43 mutant proteins that successfully dock but are non-conductive, it stands to reason that mutations precisely characterized to have altered or disrupted gating properties would likely fit the scenario in which FETCH demonstrates docking and FRAP demonstrates non-conductivity.

While complementary to existing methods, FETCH makes advancements in terms of ease of use, accessibility, and time scale. Specifically, FETCH exploits the robust expression profile of the commercially available HEK 293FT cell line; C-terminally tagged fluorescent vectors are widely-used and commercially available materials; and transient transfections avoid the costly and time consuming generation and maintenance of stable cell lines. Additionally, FETCH is directly scalable for high-throughput evaluations as flow cytometry analysis can be completed from single sample tubes or 96 well plates.

Though HEK 293FT cells are rarely reported as vehicles for connexin studies, we observed that 293FT cells express significantly greater levels of exogenously introduced Cx-FP constructs than 293 and HeLa cell lines. Moreover, though we could identify gap junction plaques in all of the cell lines we transfected, we observed substantially larger surface area of plaques in 293FTs, likely due to the comparatively elevated protein expression (Supp Figure 1). Critically, 293 and HeLa cell lines showed substantially lower FETCH scores, when expressing the otherwise robustly positive Cx43-FP constructs, thus, these cells lines were not pursued for subsequent FETCH analyses.

Here, we showed that FETCH can evaluate gap junction formation of numerous connexin proteins. Analysis of the entire complement of human connexin isoforms revealed that a majority of isoforms generate positive FETCH scores when evaluated for homotypic docking. Furthermore, when evaluating heterotypic pairings of selected isoforms representing different isoform classes, we observed positive FETCH scores in agreement with known heterotypic pairings such as Cx26/Cx32 and Cx43/Cx45, which generate heterotypic gap junctions in rodent liver and cardiomyocytes, respectively [70, 71]. Additionally, we observed a potentially novel docking interaction when pairing Cx26/Cx36, which has not been previously explicitly characterized. Finally, we demonstrated that FETCH has utility in evaluating the docking ability of disease-relevant connexin mutant proteins. Specifically, we showed that a number of ODDD-associated Cx43 mutations result in docking perturbations that were also reflected in poor fluorescence recovery in GAP-FRAP functional analyses, indicating non-functionality.

Overall, these results demonstrate that FETCH is a valuable new method that reliably evaluates connexin hemichannel docking interactions and can explore previously uncharacterized docking combinations of endogenous connexin isoforms, as well as evaluate docking features of mutant connexin proteins present in disease states. Importantly, FETCH does not quantify hemichannel binding affinity, instead, FETCH captures docking interactions in the context of several isoform specific variables (e.g., expression level, turnover rate and plaque size) that likely contribute greatly to positive score variability amongst different isoforms. Thus, FETCH scores above a defined threshold are considered an indication of docking, but variation in the scores between different isoforms are not comparable. Furthermore, FETCH scores of individual isoforms may be increased by construct and condition optimization. On the other hand, discrepancies in FETCH score between wild-type and mutant counterparts of the same connexin isoform are comparable and directly identify perturbations in docking. An important caveat is that FETCH conclusively identifies docked, compatible connexin protein interactions, however, negative FETCH scores may reflect either non-compatible counterparts or a failure of one or both proteins to localize or fold properly. Finally, FETCH does not reveal mechanism of pathology of evaluated disease-associated mutant proteins but instead sheds light on a single contributing aspect- docking ability of the mutant isoform. Thus, we believe FETCH is best utilized in conjunction with other biochemical and imaging analyses for comprehensive understanding of connexin behavior.

## METHODS

### Construct cloning and preparation

Connexin gene information was procured from the National Center for Biotechnology Information (NCBI, ncbi.nlm.nih.gov) and the Ensembl genome browser (ensembl.org). The human codon-optimized genes were ordered from Integrated DNA Technology (IDT, idtdna.com). Genes were subcloned into Emerald-N1 (addgene:53976), piRFP670-N1 (addgene: 45457) and Apple-N1 (addgene: 54567) vectors using In-Fusion cloning (takarabio.com), resulting in connexin fluorescent fusion proteins, specifically with the fluorescent proteins being adjoined to the connexin carboxy-terminus. Previous studies have indicated that carboxy-terminal tagging of connexin proteins is less likely to cause functional perturbations than amino-terminal tagging [7272].

Mutant constructs were generated using overlapping primers within standard Phusion polymerase, PCR amplifications to facilitate site-directed mutagenesis. The ‘self-cleaving’ T2A tag for multi-Cx co-expression was generated via primer extension and In-Fusion ligation. Subsequent to the generation of the T2A tag, different Cx-FP fusion genes were inserted on either side of the T2A tag, whilst maintaining all constructs inside of one open reading frame, using restriction enzymes digest and In-Fusion ligation.

### Cell Culture

HEK 293FT cells were purchased from Thermo Fisher Scientific (cat# R70007) and were maintained according to manufacturer instruction. Cultures were grown in 10cm tissue culture treated dishes in high-glucose DMEM (Sigma Aldrich, D5796) supplemented with 6mM L-glutamine, 0.1 mM MEM non-essential amino acids and 1mM MEM sodium pyruvate in a 5% CO_2_, 37°C incubator. Cells were passaged via trypsinization every 2-3 days or until 60-80% confluency was reached.

### Transient Transfection

HEK 293FT cells were plated in 10ug/ml Fibronectin coated multi-well dishes to achieve ~75% confluency after overnight incubation. For transfection, DNA was combined with polyethylenimine (PEI) diluted in Opti-MEM in a 1:3 ratio (ug of DNA: uL of PEI) and incubated at room temperature for 10 minutes. Following incubation, PEI-DNA complexes were added dropwise to wells of plated cells. Treated cells were then incubated at 37°C for 16-18 hrs, followed by media change. Expression of the connexin-FP constructs was evaluated at 24 hrs and 48 hrs post transfection via widefield or confocal microscopy and western blotting.

### Confocal Imaging Analysis

For imaging, HEK 293FT were plated in 10ug/ml Fibronectin coated 35 mm, glass-bottom Mattek dishes (cat# P35GC-1.5-14-C). Cells were transiently transfected using PEI as described above and imaged at ~48 hrs post transfection. Images were acquired on a Leica SP5 laser point scanning inverted confocal microscope using Argon/2, HeNe 594nm and HeNe633nm lasers, conventional fluorescence filters and a 63X, HCX PL APO W Corr CS, NA: 1.2, Water, DIC, WD: 0.22mm, objective. Images were taken with 1024 × 1024 pixel resolution at 200Hz frame rate.

### Cell Lysate preparation and Western Blotting

To extract cell lysates, plated cells were washed twice with room temperature PBS, trypsinized, collected and centrifuged at 2,000 × g for 5 minutes. Pellets were washed twice with PBS followed by centrifugation and aspiration. Pellets were lysed by the addition of 10 mM Tris, pH 8.0, 1 mM EDTA, 1% SDS, and 1X protease and phosphatase inhibitor cocktail (Thermo) and sonicated. Lysates were quantified via detergent compatible Bradford assay (ThermoFisher). For western blot, samples were separated using gel electrophoresis on 4-20% SDS-Page gels. Separated proteins were transferred to PVDF membrane in transfer buffer (25 mM Tris, 190 mM glycine, 20% methanol, final pH 8.3), for 1 hr at 110V. Subsequently, membranes were blocked in 5% milk for 1 hr, followed by 1hr incubation with primary antibody (1:1000) in 5% milk with shaking. Blots were then washed three times with TBST(20 mM Tris, pH 7.5, 150 mM NaCl and 0.1% Tween 20) for 5 minutes and incubated with Horse radish peroxidase conjugated secondary antibody (1:5000), followed by three washes with TBST for 5 minutes. Blots were developed using Optiblot ECL kit (Abcam) and visualized via Azure c300 phosphorimager. To restain individual blots for different proteins, blots were stripped with mild stripping buffer (200 mM glycine, 0.1% w/v SDS, 1% Tween 200), 2 × 10 minute washes with shaking, followed by 2 × 10 minute PBS washes with shaking and 2 × 5 minute TBST washes with shaking. After stripping, blots were blocked in 5% milk for 1 hr in preparation for antibody staining as previously described.

### Flow Enabled Tracking of Connexosomes in HEK 293FT cells (FETCH)

FETCH analysis is fundamentally a two-component system. To complete FETCH analysis, replica multi-well plates were transfected with either of the two components being evaluated. The media of transfected wells was changed 16-18 hrs post transfection and cells were trypsinized and experimental counterparts were combined at ~20 hrs post transfection. The entirety of combined samples were then plated onto new, 10ug/ml Fibronectin coated wells of the same size, resulting in hyperdensity and over confluency. Following co-plating, samples were incubated for ~20-24 hrs, at which point samples were trypsinized, resuspended in phosphate buffered saline with 10U/ml DNAse and fixed with paraformaldehyde (f/c of 1.5%). Co-plated samples in 96 well plates were resuspended to a final volume of ~150 ul, samples from 24-well plates were resuspended to a final volume of ~600 ul.

Flow cytometry data was collected on either a BD FACSCanto II (2-color experiments and high-throughput 96-well plates; 488nm and 633nm lasers) which utilizes the BD FACSDiVa software. Samples were analyzed in two selection gates prior to their fluorescence evaluation. First, presumable HEK cells were identified by evaluating sample forward vs side scatter area. Next, single cells were selected by evaluating cells that maintained a linear correlation of forward scatter height to forward scatter area. Finally, the fluorescence profiles of each sample was generated.

### Fetch Automated Gating Pipeline

Each FETCH experiment produces “*.fcs” files that contain all the channel data for fluorescence in the sample. Our automated pipeline loads these files, extracts FSC-A, SSC-A, and FSC-H. Depending on the machine used, we either load green channel as 1-A or as FITC-A. For the red channel, we can either have two (APC-A (RFP670) and PE-A (mApple)), or just one: 5-A(RFP670). Next, our code produces two matrices containing SSC-A with FSC-A, and FSC-A with FSC-H respectively.

Our first gate is drawn on the FSC-A vs SSC-A axes to exclude cellular debris which clusters in the lower left corner and the cells that are saturating the laser (at the max of both axes). On an FSC-A vs SSC-A plot, the cellular debris usually is smoothly transitioning into the population of intact cells, therefore we use a Gaussian kernel density estimator with the estimator bandwidth selection defined by the Scott’s Rule to draw contours around the data in SSC-A vs FSC-A matrix. We next use a set of heuristics to determine which of the contour lines should be used to define the first gate. Specifically, cellular debris usually clusters below 25000 on both axes, so any contour that includes values at or below is excluded. Similarly, any contour within a 1000 of the maximum value of each axis is also excluded. Of the remaining contours, the largest one is chosen and an oval equation is fitted to the points defining that contour to attenuate occasional protrusions of the contour that tap into cellular debris population in rare cases. The fitted oval becomes the first gate.

For all the elements inside of the first gate, a second gate is drawn in the axis of FSC-A and FSC-H to exclude non-single cells. For the second gate, first we fit a line to all of the points. Next, for each point we find a norm to the fit line and find a standard deviation of all such norms. Using this standard deviation, we define a second gate 4 standard deviations away from the fitted line on both sides and exclude all the points outside of this gate.

Upon applying the first two gates, we finally plot the data with the red fluorophore on the y axis and the green fluorophore on the x axis. If a sample contains more than two fluorophores the last gate is drawn for each possible combination. Since some readings are below zero due to fluorescence compensation, we shift all the data points by the smallest value along both axes and then take a natural log of fluorescence levels. To achieve the optimal bandwidth for the kernel density estimation, we run a cross validation grid search algorithm on the points in the log space. Then we fit a gaussian kernel density with the bandwidth estimated to obtain density contours. For properly expressing samples, we expect a large population of untransfected cells in the bottom left quadrant of the plot, a population of cells strongly-expressing red fluorophore along the y axis, and a population of cells strongly expressing green fluorophore along the x axis. We expect autofluorescence to not exceed 500 on either axis, so the untransfected population is defined to be below this value along both axes. To draw a tighter bound of the untransfected population, we choose the first contour whose mean kde value is at or above the 60th quantile (identified as a generalizable heuristic value) of the distribution of kde values within the largest contour which is at or below the autofluorescence cutoff. The top-most point of the tight contour defines the horizontal gate and the right-most point -- the vertical gate, separating the plot into four quadrants.

The upper left quadrant Q1 corresponds to the cells expressing just the red fluorophore, the upper right quadrant Q2 represents dual-colored cells, the lower right quadrant Q3 -- the cells expressing just the green fluorophore, and the lower left quadrant Q4 represents untransfected population. FETCH score is defined as the proportion of dual-colored cells to all transfected cells: Q2/(Q1 + Q2 + Q3).

Expecting approximately equal expression levels of each fluorophore, if the number of cells in Q1 is two or more times larger/smaller than the number of cells in Q3, the FETCH score is classified as “dubious” and marked accordingly in the output table. The “dubious” label is also given to samples that have less than 500 cells total after the application of the second gate and to the samples that failed at any of the steps in the pipeline (usually due to poor expression or the absence of cells in the sample).

### Fluorescence Recovery after Photobleaching (FRAP) Analysis

For FRAP analysis, HEK 293FT cells were plated into 10ug/ml Fibronectin coated mattek glass-bottom dishes (MatTek Corporation, cat#: P35G-1.5-14-C) to achieve ~75% confluency after overnight incubation. Dishes were then transfected with connexin constructs and allowed to incubate overnight. After 16-18hrs, transfection media was replaced with fresh media and FRAP was initiated at ~24 hrs post transfection. For FRAP, 1ug/ml Calcein-AM (Enzo Life Sciences, cat#: 52002) was added to dishes, incubated at 39°C, 5% CO2 for 30 minutes and then aspirated. Dishes were rinsed 3 times with phosphate buffered saline and then imaged in Tyrode’s Solution (Alfa Aesar, cat#:J67593-AP).

Imaging was completed on a Leica SP5 confocal microscope using the LAS Software FRAP Wizard. The experimental program centered on imaging with a 63X water immersion objective, 512 × 512 Hz and staging pre-bleach (10 frames, 1.29s/f), bleach (30 frames, 1.29 s/f) and recovery (24 frames, 10s/f) imaging intervals. For bleaching conditions, the Argon laser was set to 20% power and full power was used to bleach a selected cell ROI. Cells expressing the connexin construct of interest were selected for bleaching if they were surrounded by at least 3 cells making clear contacts and also clearly expressing the exogenous construct, as determined by fluorescence.

FRAP images were analyzed using a custom written FIJI plugin and python script. Using a semi-automatic FIJI plugin, 2 ROIs were drawn: one around the bleached cell, and the other -- around a cell that did not get bleached. Raw fluorescence was extracted from those and saved into a file for each frame in a stack. Files were then processed using the custom python script, first normalizing the fluorescence of the bleached cell to the fluorescence of the unbleached cell for each frame, and then calculating the percent fluorescence of the baseline by dividing the resultant value by the mean corrected value of the pre-bleaching frames. Since the majority of mutants exhibited low to no fluorescence recovery, it was impossible to fit a reasonable exponential curve to them, impeding the calculation of time constant and similar fitted metrics. Instead, we used the percent of the baseline value at the end of an imaging experiment for statistical analysis.

---

**Supplemental Figure 1:**
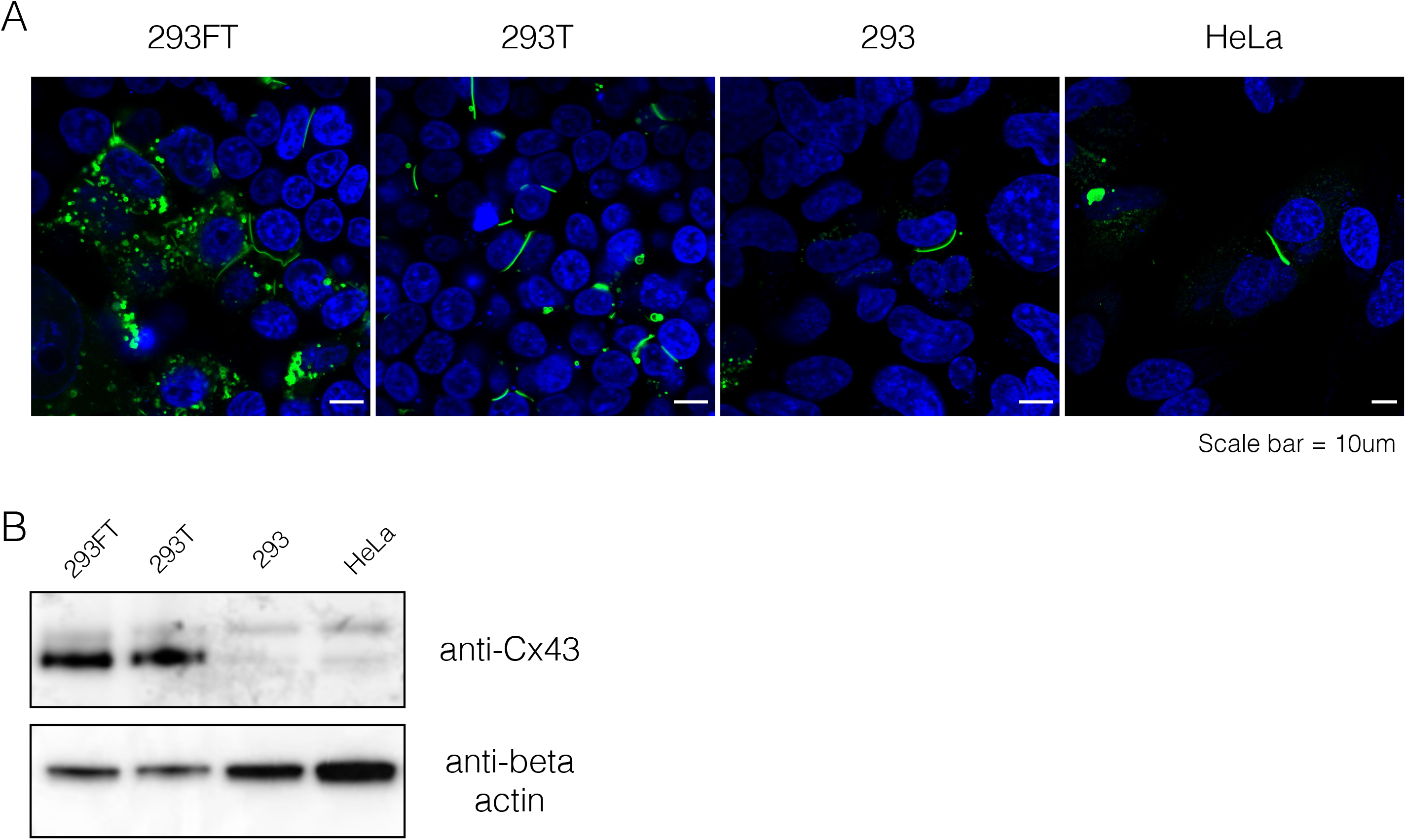
Expression of Cx43-mEmerald in different cell lines. (A) Confocal images of 293FT, 293T, 293 and HeLa cell lines transiently transfected with Cx43-mEmerald, nuclei are stained with DAPI. (B) Western blot of different cell lines transiently transfected with Cx43-mEmerald.

**Supplemental Figure 2:**
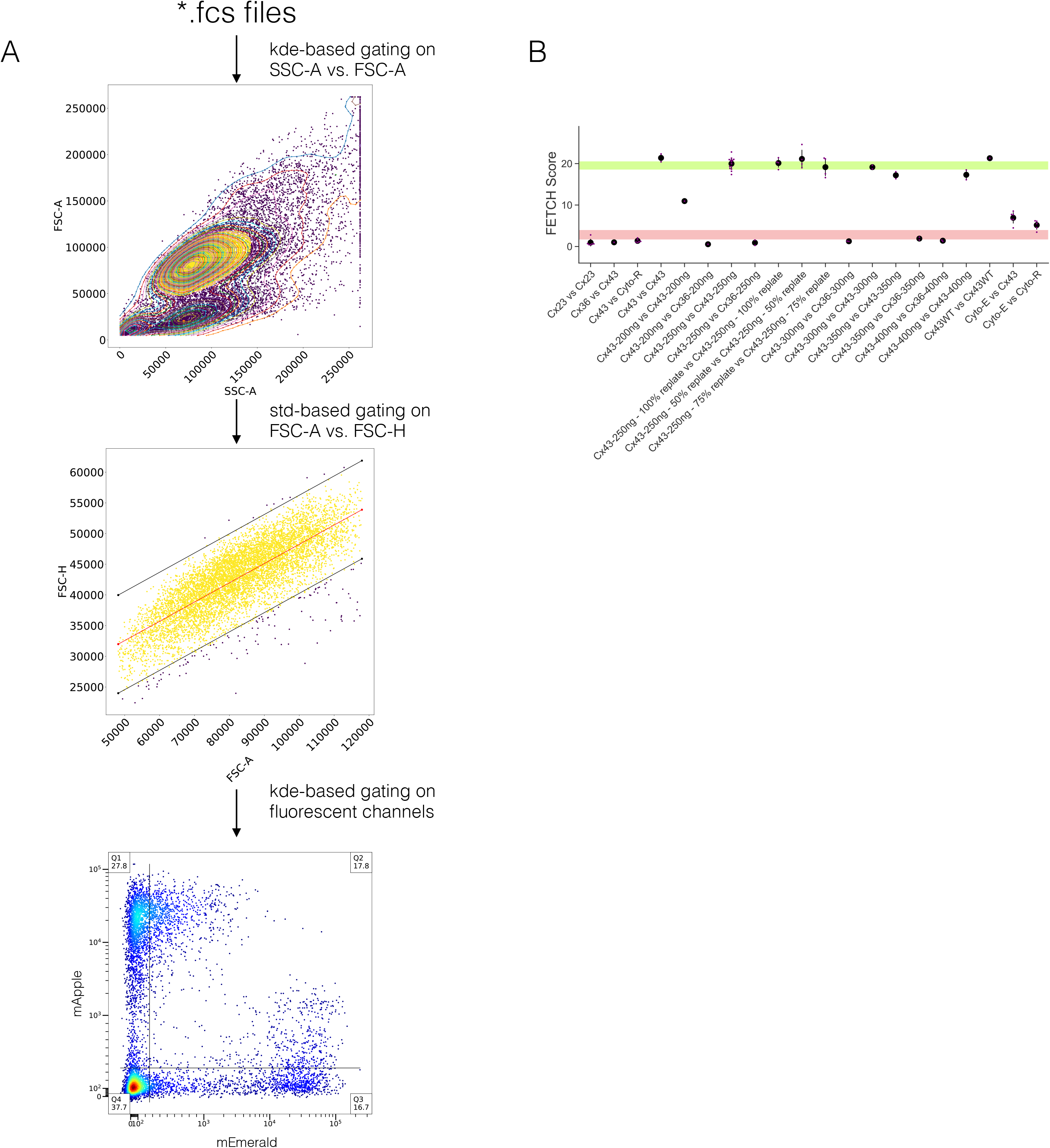
FETCH analysis pipeline and pilot data for FETCH score distributions. (A) Automated gating pipeline that uses sequential kde- and std-based gating approaches to obtain a FETCH score from a single raw .fcs file. (B) FETCH score distributions of known positive and negative combinations used for statistical evaluation. The 95% confidence interval for the known positive and known negative distributions are shown in green and red, respectively.

## AUTHOR CONTRIBUTIONS

Conceptualization and Methodology, E.R.; Formal Analysis, E.R., K.C., and K.D.; Investigation, E.R.; Resources, N.B.; Writing -Original Draft, E.R., K.D., Writing – Review & Editing, E.R., K.C., N.B., and K.D.; Visualization, E.R., K.C.; Supervision, N.B and K.D.; Project Administration, and Funding Acquisition, E.R., N.B., and K.D.

## ACKNOWLEDGEMENTS

The authors would like to acknowledge Dr Terry Oas for helpful discussions and guidance and Dr. Mike Cook for flow cytometry analysis support. The authors declare no competing financial interests. The authors would like to acknowledge the generous funding that has supported this work: Duke University School of Medicine MedX Grant (K.D. and N.B.), Duke University Chancellor’s Discovery award (K.D.), NIH grant R21EY029451 (K.D), Duke University School of Medicine Kahn Neurotechnology Grant (K.D.), Ernest E. Just Life Science Institute Postdoctoral Research Fellowship (E.R.), and NIH grants U01HL134764, R01HL132389, and R01HL126524 (N.B.)

